# Emergent honeycomb topology of the leaf spongy mesophyll

**DOI:** 10.1101/852459

**Authors:** Aleca M. Borsuk, Adam B. Roddy, Guillaume Théroux-Rancourt, Craig R. Brodersen

## Abstract

The spongy mesophyll layer in leaves is ubiquitous among vascular plants, yet its structure is relatively unknown and typically described as a disordered assemblage of isodiametric cells. We characterized spongy mesophyll structure among diverse taxa using X-ray microCT imaging and found that leaves with small cell sizes, high cell packing densities, and close vein spacing were congruent with the isodiametric paradigm. When these structural traits exceeded well-defined thresholds, the spongy mesophyll domain was instead tessellated with an emergent topological motif of an irregular honeycomb that minimizes cellular investment and obeys Euler’s Law of space filling. Our data suggest spongy mesophyll is governed by allometric scaling laws, with two distinct topologies optimized for either photosynthetic performance or minimal resource investment.

**One Sentence Summary:** Conserved topological motifs in the spongy mesophyll are coordinated with leaf photosynthetic performance.

## Main Text

The laminar leaf with reticulate venation is common among terrestrial vascular plants, and convergence on this form has occurred independently in at least four distinct lineages since the Paleozoic (*1*). Despite large variation in leaf size, shape, and structure (*2*), the interior of most laminar leaves has a conserved morphological scheme consisting of two photosynthetic layers arising from a common dorsiventral developmental framework (*3*): the palisade and the spongy mesophyll. The palisade mesophyll is generally positioned immediately below the upper epidermis and is composed of cylindrically shaped cells oriented perpendicular to the leaf surface. This layer is characterized by a high surface area to volume ratio that facilitates CO_2_ absorption in a region of the leaf where light is abundant and photosynthetic rates are high (*4*–*7*). The spongy mesophyll, positioned below the palisade, is generally treated as a disordered and loosely packed assemblage of isodiametric, or roughly spherical, cells. It is thought that this tissue satisfies multiple biophysical roles (*8*) and promotes both light scattering and CO_2_ conductance from the stomata to the palisade (*9*, *10*). However, testing such hypotheses requires detailed structural characterization that has been difficult to achieve due to the scale, complexity, and three-dimensional (3D) organization of the spongy mesophyll (*11*). Although specific examples of non-isodiametric cell shapes in the spongy mesophyll have been reported (*8*, *12*– *15*), limited data exist on the structural organization of this tissue or how structural variation influences leaf function. We sought to determine the 3D geometry of the spongy mesophyll using X-ray microCT imaging to investigate the variation in spongy mesophyll architecture, and to explore relationships between mesophyll structural properties and photosynthetic traits.

### Structural characterization

Using X-ray micro-computed tomography (microCT) imaging (*5*, *11*, *16*) we extracted 3D anatomical traits of the spongy mesophyll for 40 species with laminar leaves and reticulate venation that represented every major clade of terrestrial vascular plants, with several congeneric pairs for deeper investigations into the extent of variation between closely related species (Data Set S1). This analysis revealed two dominant spongy mesophyll classes (Fig. 1) with low topological diversity across the sampled species. Spongy mesophyll class was defined by its organization in the paradermal plane, i.e. parallel to the leaf surface. In the first class (27% of the species), spongy mesophyll conformed to the established isodiametric model, and was composed of a network of interconnected small-diameter cells surrounded by continuous intercellular airspace (IAS; Fig. 1A,B; *16*). By observation, this class of spongy mesophyll has a disordered 3D network best described as an open-cell foam (*17*).

**Fig. 1.**
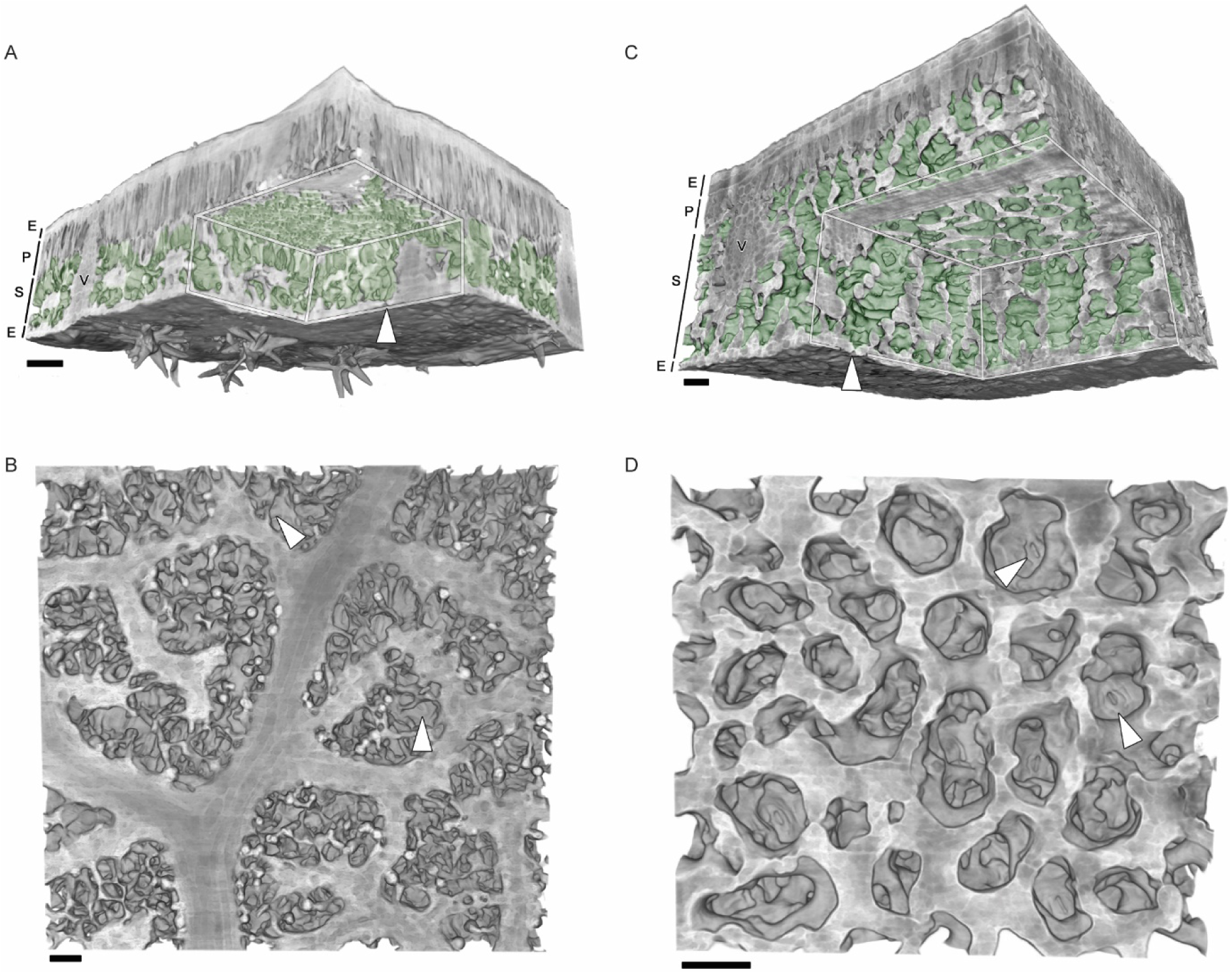
3D views of the two primary topological classes of spongy mesophyll imaged with microCT. **(A)** Representative leaf with the non-honeycomb spongy mesophyll (*Quercus suber*) with cropped interior showing intercellular airspaces (green) within an irregular network of spongy mesophyll (S). Epidermis (E), palisade mesophyll (P), and vascular tissue (V) shown in grayscale. Stomate indicated by the white arrow. Scale bar = 50 µm. **(B)** Paradermal view of the same *Q. suber* leaf’s spongy mesophyll and veins. **(C)** Representative leaf with the honeycomb topology (*Berberis nervosa*) with cropped interior showing columnar intercellular airspaces (green) within the spongy mesophyll (S). The position of a representative stomate is indicated with a white arrow, showing spatial coordination with an adjacent column of intercellular airspace. Epidermis (E), palisade mesophyll (P), and vascular tissue (V) shown in grayscale. Scale bar = 50 µm. **(D)** Paradermal view of the same *B. nervosa* leaf’s spongy mesophyll.

In contrast, spongy mesophyll in the second class was composed of layers of mesh-like tissue that conformed to the topological principles governing a honeycomb (Fig. 1C,D; *17*). The honeycomb topology was present in 72.5% of the species sampled (21 of 30 genera). Congeneric sampling revealed that spongy mesophyll topology was conserved within the genus for our dataset (Data Set S1). While we restricted our analysis to the former two groups for species with reticulate venation, we note that deviation from those two classes occurred, including in spinach (*Spinacia oleracea*, Fig. S1A) with highly elongated cells forming large and continuous airspace volumes, and in the aquatic water lily, *Nuphar polysepela*, with elongated chambers presumably to increase the air-space volume required for buoyancy (Fig. S1B). A distinctive third class of spongy mesophyll was observed in leaves with parallel venation (n = 4), in which strands of elongated mesophyll attached to veins at right angles, forming an approximately rectilinear lattice (Fig. S2).

Honeycomb mesophyll was composed of vertically stacked and horizontally aligned lattice layers. IAS columns formed by this alignment were typically positioned above stomata, forming conduits through the vertical profile of the leaf (Fig. 1C; Movies S1, S2). This supports the concept, suggested as early as 1914 by Haberlandt (*8*), that a primary function of the spongy mesophyll is to maximize CO_2_ diffusion from stomata to the palisade, and provides insight into the shape and size of the substomatal cavity (*18*, *19*). However, IAS connectivity was not strictly unidirectional. Rather, the vertical alignment of the individual mesophyll cell layers was often slightly offset or contained gaps, which would allow for lateral diffusion of CO_2_ between adjacent IAS columns (*20*, *21*).

To quantitatively examine the structure of the honeycomb spongy mesophyll (n = 29 species), we used the framework established by Gibson and Ashby (1999) for cellular solids (Data Set S1), for which honeycomb properties are derived from the packing scheme in the principal plane, i.e. the paradermal plane for plant leaves. Traits (Data Set S1) were measured on paradermal slices of spongy mesophyll (Fig. 2A,B) obtained via microCT imaging (Fig. 2C) and validated with fluorescence microscopy (Fig. 2D). At the cellular scale, the honeycomb structure was composed of cells with arm-like protrusions with a variety of individual morphologies (Fig. 2D). Yet, the tissue scale organization was largely invariant, highly ordered, and obeyed the topological constraints of Euler’s Law (*17*), which governs how geometrical indices give rise to aggregate properties for continuous 2D honeycombs, and predicts the formation of hexagons from triply-joined vertices of a lattice. The most efficient honeycombs are hexagonal lattices, which minimize investment in materials needed to tessellate a 2D plane (*22*). Hexagonal honeycombs are characterized by their vertices that sit at the junction where edges meet, with a mean edge connectivity (*Z*_*e*_) of three, a mean number of edges 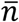 surrounding the hexagons of six, and a mean internal angle between edges (θ) of 120° (Fig. 2E). Among species within our dataset, the spongy mesophyll tissue lattice had vertices that were joined by *Z*_*e*_ = 3.03 ± 0.02, the IAS was partitioned into polygons with 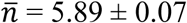, and the characteristic θ of the IAS polygons was 118.86° ± 0.40° (Fig. 2F, Data Set S1). To quantify the degree of order in the structure, we calculated the tessellation entropy (*S*) for each sample, which would be zero for a perfectly ordered hexagonal honeycomb. We found a mean tessellation entropy of 1.43 ± 0.03, which is similar to values reported for irregular hexagonal honeycomb morphologies in engineered thin films (*S* = 1.48; *23*). We also found evidence for a distribution of IAS polygon sizes and classes (Fig. 2G,H) within the void space of the honeycomb lattice that satisfied additional topological rules anticipated from a contiguous 2D honeycomb lattice. As given by the Aboav-Weaire Law (*24*) for neighboring polygon relationships, the occurrence of an IAS polygon with a lower than average number of edges (n < 6) introduced a corresponding polygon with a higher number of edges into the aggregate (Fig. 2I, upper panel). As expected from Lewis’ Rule of polygon size dispersion (Fig. 2I, lower panel), which is derived from the space filling properties of plant epithelial cells (*25*), the area of a given IAS polygon varied linearly with its number of edges. These data indicate that the spongy mesophyll and IAS matrix meet the assumptions for a hexagonal honeycomb with some irregularity as would be expected in a biological system (*17*) where multiple stresses act upon the structure throughout development. Taken together, these indices (Fig. 2F-I) show the applicability of the 2D honeycomb framework to spongy mesophyll tissue and quantify the dispersion of sizes and shapes of the IAS voids, which may provide insight for future work into the developmental processes by which the tissue was formed (*17*). We conclude that an irregular hexagonal honeycomb topology characterizes a common spongy mesophyll phenotype that spans a wide range of terrestrial plants, including representatives from the fern, gymnosperm, basal angiosperm (ANITA), magnoliid, and eudicot clades. The observation that some cells within the honeycomb lattice can be isodiametric while others have varying numbers of arm-like protrusions (Fig. 2D) implies that this structure may be formed by an arbitrary number of configurations of cell morphologies that, at the tissue scale, result in conservation of the global, ordered honeycomb topology governed by simple rules of cellular organization and optimality of resource allocation. Our data suggest that leaves with the honeycomb topology have optimized the construction of the spongy mesophyll to minimize cellular investment in a way that facilitates the vertical diffusion of CO_2_ from the stomatal pore, through the spongy mesophyll layer, and into the palisade tissue where light is most abundant for photosynthesis. This finding weakens the paradigm of spongy mesophyll as inherently disordered, and suggests that the emergent tissue structure is driven by conserved developmental (*26*), biomechanical, or physiological requirements, with distinct spongy mesophyll classes optimized for divergent functional goals.

**Fig. 2.**
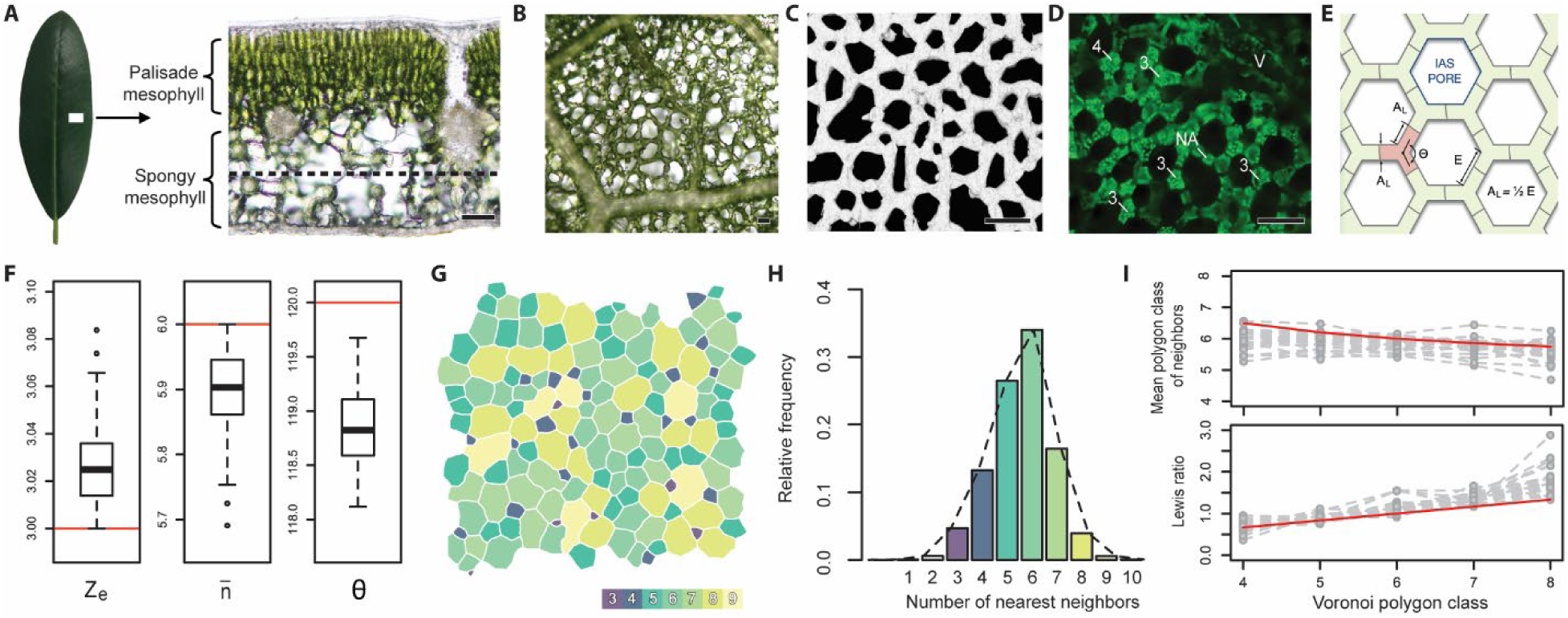
Irregular honeycomb topology and structural characterization. **(A)** Cross-section of a representative leaf (*Rhododendron* sp. shown in panels A-D; scale bars = 50 µm). **(B)** Paradermal section showing spongy mesophyll between veins. **(C)** Paradermal microCT image of spongy mesophyll layer. **(D)** Fluorescence microscopy showing spongy mesophyll cell walls (dark lines) and chloroplasts (green points). Cell arm counts (3, 4, NA) and vascular tissue (V) indicated. **(E)** Schematic diagram of honeycomb structure with cell walls (gray) outlining spongy mesophyll (tissue in green, individual cell in red) with arm length (A_L_) and arm diameter (A_D_). Arms form edges (E) that enclose intercellular airspace pores (IAS Pore) with internal angles (θ). **(F)** Box plots for mean edge connectivity (*Z*_*e*_), mean edges per face (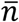), and mean internal angle (θ) for n = 29 species with the honeycomb phenotype. Boxes represent inter-quartile range, lines across boxes represent group median, and whiskers extend from the upper and lower quartiles to the group maximum and minimum, respectively. Asterisks represent sample outliers. Red bars indicate values for a perfect hexagonal honeycomb. **(G)** Nearest neighbor diagram for a representative sample. **(H)** Frequency distribution of nearest neighbors for species with the honeycomb phenotype. **(I)** Comparison of predicted (red) and measured (gray) values for dispersion in polygon properties for the Aboav-Weaire Law (upper panel) and Lewis’ Rule (lower panel).

### Drivers of phenotypic variation

To understand what factors were associated with variation in spongy mesophyll phenotype, random forest analysis was used to rank the importance of various anatomical, environmental, and taxonomic traits (Data Set S1) in classifying mesophyll as honeycomb (discretized IAS domains in the paradermal plane, n = 29; Fig. 1C,D) or non-honeycomb (continuous IAS in the paradermal plane, n = 11; Fig. 1A,B). Spongy mesophyll phenotype was best predicted by three anatomical traits (Fig. 3A): cell arm length (A_L_), cell packing density, and the characteristic minimum vein spacing (see Supplementary Materials). Mesophyll classification showed steep transitional thresholds from honeycomb to non-honeycomb phenotypes at A_L_ values below ~12 µm (Fig. 3B), as cell packing density exceeded ~2000 cells mm^−2^ (Fig. 3C), and as minimum vein spacing fell below ~0.1 mm (Fig. 3D). Therefore, leaves with smaller, more densely packed spongy mesophyll cells and closely spaced veins were more likely to exhibit non-honeycomb topology with continuous IAS domains, while leaves with fewer, larger cells per unit leaf area and more distantly spaced veins were more likely to exhibit the honeycomb topology with columnar airspace domains. The transition between phenotypes also corresponded to notable thresholds (Fig. S5) in vein density (D_v_) and stomatal density (D_s_). The non-honeycomb phenotype was associated with D_v_ above ~9 mm mm^−2^ and D_s_ above ~260stomata mm^−2^, which are trait values typically associated with angiosperm eudicots (*27*) and correlated with high photosynthetic rates (*28*). Cell arm length was significantly correlated (Spearman correlation, r_s_) with other leaf traits (Table S1), such that increases in cell arm length generally reflected increases in the characteristic dimensions of the entire structure, e.g. cell arm diameter (r_s_ = 0.95, p < 0.001), stomatal guard cell length (r_s_ = 0.78, p < 0.001), and the diameter of the IAS voids in the honeycomb lattice (r_s_ = 0.83, p < 0.001). These relationships retained significance after accounting for shared evolutionary history (Table S2).

**Fig. 3.**
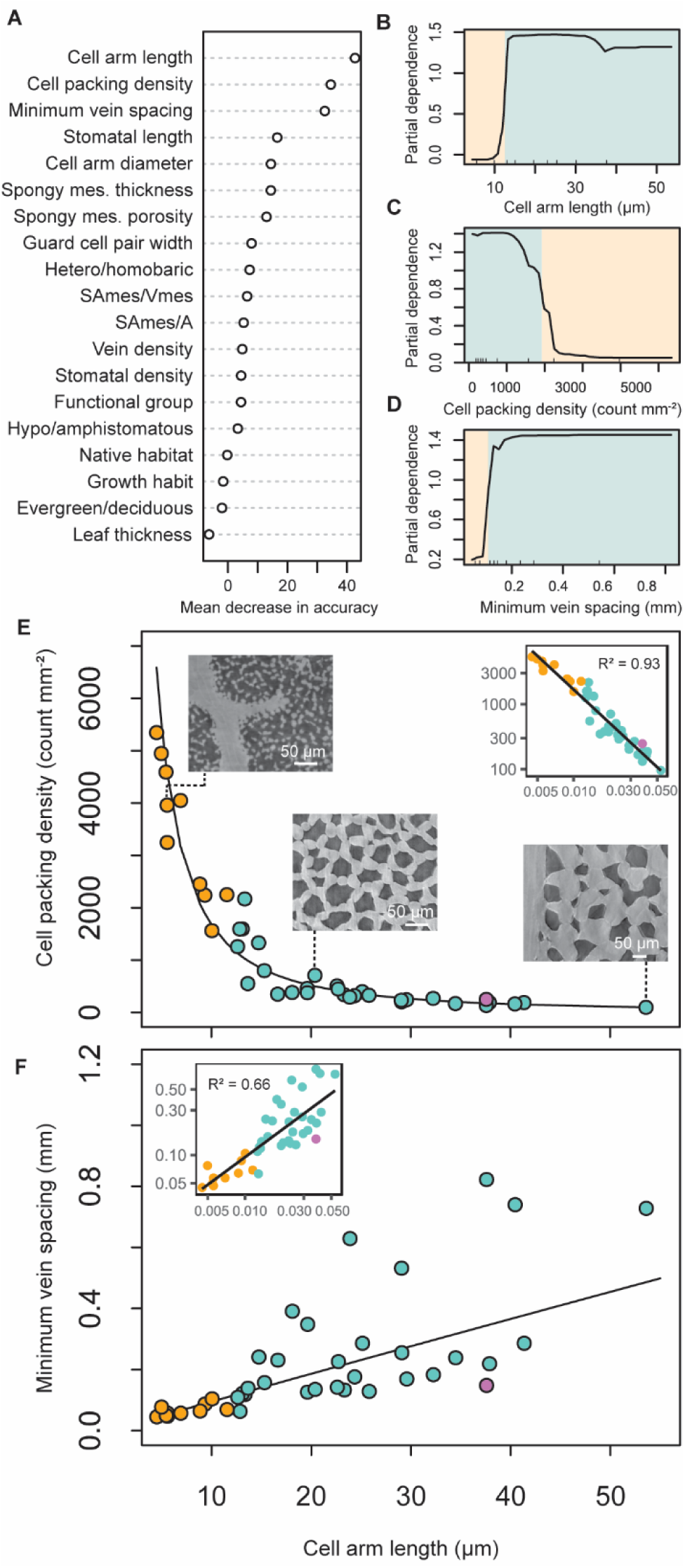
Random forest classification of spongy mesophyll phenotypes. **(A)** Variable importance for classification between honeycomb and non-honeycomb phenotypes. Predictors at the top of the ranking have a higher relative importance in classification, as determined by mean decrease in accuracy when these variables are removed from the model. **(B-D)** Partial dependence plots showing logits (log fraction of votes) as a function of the three traits with highest relative importance. Blue shading shows approximate values over which probability favors the honeycomb phenotype; orange shading shows approximate values over which probability favors the non-honeycomb phenotype. **(E)** Power law relationship (solid line) between A_L_ and cell packing density; species with honeycomb and non-honeycomb spongy mesophyll are shown with blue and orange circles, respectively. *S. oleracea*, which was treated as non-honeycomb in the analysis yet had a unique phenotype, is indicated with a purple circle. Image insets demonstrate spongy mesophyll variation throughout the trait space. Figure inset shows the log-log transformed data and linear fit (solid line). **(F)** Power law relationship (solid line) between A_L_ and minimum vein spacing. Color scheme and figure inset are the same as in (E).

With the indication that cell size, cell packing density, and vein spacing played an important role in phenotypic variation, we then explored the relationships between these traits. First, cell packing density followed power law scaling with A_L_ (R^2^ = 0.93, F(1,38) = 520.2, P < 0.001; Fig. 3E), showing that increasing that cell packing density resulted from shortening cell arm length. We note that cell counts taken from the transverse plane (as is typically done with light microscopy; *16*) could lead to overestimations of cell packing density because cells with more articulated or more numerous arms could be counted multiple times, confounding the patterns in space filling and cellular organization apparent in the paradermal plane. Cell packing density was lower in species with long cell arms, which may reflect either a lower number of cell divisions or larger minimum cell size in the spongy mesophyll during development (*29*).

Minimum vein spacing and A_L_ scaled allometrically (R^2^ = 0.66, F(1,38) = 74, P < 0.001, Fig. 3F), with eudicots having both the shortest distances between veins and the smallest cells. This relationship between minimum vein spacing and cell size is consistent with previous work showing that early Cretaceous angiosperms underwent reductions in cell size and increases in cell packing densities that elevated the conductance to CO_2_ and water necessary to increase photosynthetic rates (*27*, *28*, *30*–*32*). In terms of biomechanics, leaves with higher vein density have an abundance of semi-rigid xylem conduits that provide mechanical support (*27*). In contrast, the honeycomb structure itself is positioned between the upper and lower epidermises like in a sandwich beam (*33*), and could produce a lightweight material that is elastic when loaded in the paradermal plane and stiff when loaded normal to the leaf surface, much like the manufactured honeycombs used in packing materials (*17*).

### Structural trait scaling and functional implications

Cells of the spongy mesophyll were best characterized by a power law relationship between the length and diameter of the cell arms (R^2^ = 0.81, F(1,35) = 148.3, P < 0.001; Fig. 4A) with the honeycomb phenotype represented in gymnosperm (Fig. 4B), fern (Fig. 4C), ANITA (Fig. 4D), magnoliid (Fig. 4E), and eudicot (Fig. 4F) species, and the non-honeycomb phenotype represented only in eudicots (Fig. 4G). There were two notably unoccupied sectors of this two-dimensional trait space. First, no species in our dataset were found with large, isodiametric cells. Such a cell would have an extremely large volume, as found in succulent and epiphytic plants (*16*, *34*). Such species may operate with a different biochemical pathway (CAM vs. C3), where large cell volume favors malic acid accumulation and water storage, as opposed to optimization of the mesophyll for high rates of liquid and vapor diffusion necessary for C3 photosynthesis (*31*). Second, there were no species that had cells with highly elongated, narrow arms. This may be due to biomechanical limitations, or to conserve a minimum distance between the nuclear envelope and the cell membrane for intracellular translocation.

**Fig. 4.**
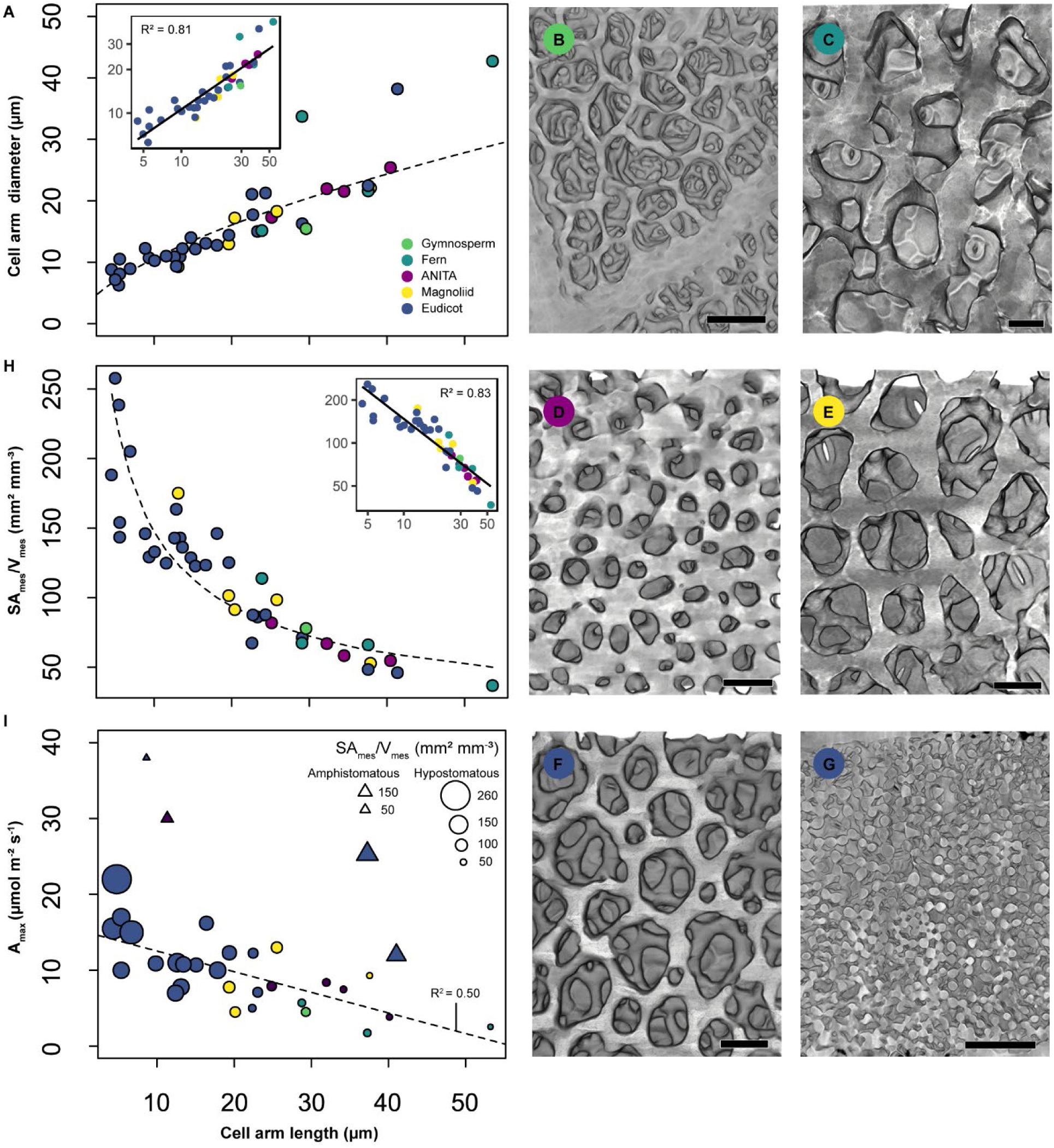
Power law scaling of spongy mesophyll cell dimensions and models of derived photosynthetic properties. **(A)** Relationship between spongy mesophyll cell arm diameter (A_D_) and arm length (A_L_). Power law regression shown by the black dashed line. Inset shows log-log transformed data and linear fit (solid line). Colors represent sample phylogenetic affinity. **(B-G)** Spongy mesophyll structure viewed from the paradermal plane for representative species from gymnosperm (*Gnetum gnemon*, B), (continued on next page) fern (*Platycerium andinum*, C), ANITA angiosperm (*Amborella trichopoda*, D), magnoliid (*Calycanthus occidentalis*, E), and eudicot (*Coffea arabica*, F; *Helianthus annuus*, G) clades. Scale bars = 50 μm. **(H)** Relationship between spongy mesophyll surface area to volume ratio (SA_mes_/V_mes_) and A_L_. Power law regression shown by the black dashed line. Inset and colors scheme are the same as in **(A)**. **(I)** Linear relationship between A_max_ and A_L_. Open circles represent hypostomatous species. Open triangles represent amphistomatous species, which were excluded from the regression. Point size scaled according to SA_mes_/V_mes_, which is linearly related to A_max_ (R^2^ = 0.62). Colors represent sample phylogenetic affinity.

To explore how spongy mesophyll structural traits predict photosynthetic properties of the leaf, we first modeled the spongy mesophyll cell surface area per unit tissue volume exposed to the intercellular airspace (SA_mes_/V_mes_) as a function of cell arm length. SA_mes_/V_mes_ is a measure of the evaporative and absorptive surface exposed to the IAS, and represents an intermediate step in the liquid-vapor pathway between veins and stomata. SA_mes_/V_mes_ is, therefore, a critical link between vein density and stomatal conductance in optimizing the hydraulic and diffusive pathways to increase leaf photosynthetic capacity (*35*). We found that as arm length increased, SA_mes_/V_mes_ of the spongy mesophyll sharply decreased according to a power law (R^2^ = 0.83, F(1,38) = 188.7, P < 0.001; Fig. 4H). Thus, plants with larger cells such as the fern *Platycerium andinum* (Fig. 4C), which were also more likely to exhibit the honeycomb phenotype, had lower SA_mes_/V_mes_ in the spongy mesophyll compared to eudicots, such as *Helianthus annuus* (sunflower), with smaller cells and the non-honeycomb phenotype (Fig. 4G). Although cell size strongly influenced SA_mes_/V_mes_, variation in cell geometry may also play a role (*14*, *15*). To demonstrate the implications of cell shape in our dataset, we calculated the surface area-to-volume ratio (SA/V) for idealized geometrical models of isodiametric and triply-armed cells using measured cell arm lengths and diameters to parameterize the analysis (Fig. S6). As anticipated, surface area increased in triply-armed cells compared with isodiametric cells of the same volume, with a mean difference in SA/V between the two modeled cellular geometries of 0.069 µm^2^ µm^−3^ (s.d. = 0.029). Thus, in addition to cell size and cell packing density, cell shape can regulate SA_mes_/V_mes_.

Given that spongy mesophyll structure and surface area provide the physical basis for the conductance of CO_2_ for photosynthesis, we lastly tested how leaf-level maximum photosynthetic rate (A_max_) was related to A_L_ and SA_mes_/V_mes_ of the spongy mesophyll (Fig. 4I). A_max_ decreased linearly with increasing A_L_ (R^2^ = 0.50, F(1,27) = 26.65, P < 0.001; Fig. 4I). The highest A_max_ values occurred among species with non-honeycomb spongy mesophyll (Data Set S1), indicating a shift from lower to higher photosynthetic capacities as the leaf undergoes structural changes from honeycomb to non-honeycomb phenotypes. Species with amphistomatous leaves such as *H. annuus* and *Gossypium hirsutum* (cotton) were excluded from the model (shown in in Fig. 4I as open triangles), as the capacity for gas exchange on both sides of the leaf promotes higher photosynthetic rates (*36*). A_max_ increased linearly with increasing spongy mesophyll SA_mes_/V_mes_ (R^2^ = 0.63, F(1,27) = 46.06, P < 0.001); thus, although the palisade mesophyll is typically modeled with a higher photosynthetic capacity relative to the spongy mesophyll (*7*), there is a strong positive relationship between the quantity of photosynthetically active surface area per unit volume in the spongy mesophyll and leaf-level photosynthetic capacity.

## Discussion

Given the historical dominance of 2D transverse analysis of leaves (Fig. 2A), our data highlight the importance of 3D characterization of mesophyll structure (*5*, *15*), scaling relationships between morphological variation and tissue mass (*37*), and how tissue geometry influences the leaf economics spectrum (*38*). Investment in increased vein density and stomatal density enabled elevated rates of photosynthesis among the angiosperms, and these traits are apparently coordinated with a striking structural shift from the irregular honeycomb topology to the non-honeycomb topology. Our data suggest that cell size and cell packing are critical for the development of a spongy mesophyll structure optimized for high SA_mes_/V_mes_ and the absorptive surface for CO_2_ acquisition (Fig. 4). Because cell size and cell packing density are fundamentally limited by genome size (*32*, *39*), streamlining the genome makes possible the miniaturization of xylem conduits, stomatal guard cells, and mesophyll cells, which collectively allow for higher photosynthetic capacity by optimizing the hydraulic and diffusive pathways in the leaf (*30*–*32*).

The widespread occurrence of the honeycomb topology suggests selection has favored minimizing cellular investment deep in the leaf where photosynthetic cells are often light-limited. Honeycombs have been found widely in natural and engineered systems as multifunctional materials for fluid transport, energy conversion, and structural support (*40*). These patterns have been observed within different types of plant tissues, from epithelial cells (*25*) to the venation pattern of reticulate leaves (*41*) where a hexagonal honeycomb topology optimizes transport efficiency of the vascular system (*42*). Based on our dataset of dorsiventral leaves with reticulate venation, it also appears that the honeycomb form may be self-similar, where the spongy mesophyll repeats the pattern of the vasculature at a smaller scale. Thus, the honeycombs appear in multiple tissues and across scales within the leaf. It is possible that the 3D honeycomb structure is driven by the physical processes and stresses that arise during development rather than functional constraints alone. Temporal and spatial coordination has been observed, for example, between mesophyll airspace and epidermal cell differentiation during development (*43*). The hexagonal tessellation of the spongy mesophyll domain may be the most efficient means of meeting multiple functional demands within a single tissue type; i.e. moving water over long distances outside the xylem, maintaining high diffusive conductance to CO_2_, exporting the products of photosynthesis, and serving as a self-supporting structure when vein density is low. Hence, for plants with relatively large mesophyll cells, distantly spaced leaf veins, and moderate to low photosynthetic capacities, the emergent topological properties of the irregular hexagonal honeycomb structure provide an alternative strategy for resource allocation, a key trait dimension across the global spectrum of plant form (*44*). This work provides a new framework for exploring leaf form-function relationships, developmental processes, and physiological models (*11*).

## Supporting information

Data Set S1

## Acknowledgments

We thank Professor Lorna Gibson (Massachusetts Institute of Technology) for discussion, and Professor Tim Brodribb (University of Tasmania) and Professor Erika Edwards (Yale University) for providing feedback on a draft manuscript. We additionally thank the Marsh Botanical Garden (New Haven, CT), the University of California Botanical Garden (Berkeley, CA), the UC Davis Botanical Conservatory (Davis, CA), and the UC Davis Arboretum (Davis, CA) for plant material, and Kyra Prats (Yale University) for assistance with maximum photosynthetic rate data collection. GTR acknowledges the Paul Scherrer Institut, Villigen, Switzerland for provision of synchrotron radiation beamtime at beamline TOMCAT.

## Author contributions

A.M.B. conceived of the original research plan and jointly with C.R.B. designed the methods, acquired, analyzed, and interpreted the data, and drafted and revised the manuscript. A.B.R. acquired leaf microCT data and contributed to data analysis and manuscript revision. G.T.R. acquired leaf microCT data and contributed to manuscript revision.

## Data and materials availability

All data are available in the manuscript or the supplementary materials.

## Supplementary Materials

Materials and Methods

Figures S1-S6

Tables S1-S3

Movies S1-S2

References (*45-87*)

