## Supplemental Material for "Emergent honeycomb topology of the leaf spongy mesophyll"

**This PDF file includes:**

Materials and Methods
Figs. S1 to S6
Tables S1 to S3
Captions for Movies S1 to S2

**Other Supplementary Materials for this manuscript include the following:**

Data Set S1
Movies S1 to S2

### Materials and Methods

Plant material

Mature, fully expanded leaves were selected from various glasshouses and arboreta and represent a broad diversity of plant groups, species native habitats, and leaf structure (Data Set S1 for full names and growth location of the species). Only laminar leaves were selected for this study, which facilitated the comparison of structural properties in the spongy mesophyll. Such a comparison would have been less straightforward with needles, scale-like, or succulent leaves with highly divergent geometries. Healthy, well-watered plants were selected, and the petiole or stem was excised, wrapped in wet paper towels, and immediately put in plastic bags. They were then transported to the MicroCT facility and scanned within 36 h of collection.

MicroCT data acquisition and reconstruction

MicroCT data acquisition was carried out at the Advanced Light Sources (ALS) at the Lawrence Berkeley National Lab (Berkeley, CA) and at the TOMCAT Tomography beamline, Swiss Light Source (Paul Scherrer Institute, Villigen, Switzerland). Reconstruction methods followed those described by Théroux-Rancourt et al. (2017). Samples were prepared before each scan (less than 30 min), by excising a ∼1.5- to 2-mm-wide and ∼15-mm-long piece of leaf tissue near approximately 33 to 50% of the distance from the leaf tip (i.e. approximately the mid point of the leaf) and offset approximately 5-10 mm from the edge of the mid vein. The cut edges of the tissue samples were oriented parallel to the nearest second order vein if possible to capture areoles bounded by high vein orders (i.e. greater than or equal to second order veins, depending on the species). Tissue samples were then enclosed between two pieces of Kapton (polyimide) tape to prevent desiccation while allowing high X-ray transmittance. They were mounted in the sample holder, centered in the microCT X-ray beam, and scanned using the continuous tomography mode capturing 1,025 (ALS) or 1800 (SLS) projection images at 21 keV, using a 5x, 10x, or 40x objective lens, yielding a final pixel resolution between 1.277 - 0.1625 µm. Each scan was completed in 5 min (SLS) or ∼15 min (ALS).

Image reconstruction was carried out using TomoPy, a Python-based framework for reconstructing tomographic data, for all ALS samples, or using the in-house reconstruction platform for SLS samples. Each raw data set was reconstructed using both the gridrec and phase retrieval reconstruction methods. Image stacks were cropped to remove tissue that was dehydrated, damaged, or contained artifacts from the imaging or reconstruction steps. The final stacks contained ∼500-2000 eight-bit grayscale images (downsampled from 32-bit images). Image processing was applied equally among scans using the FIJI distribution of ImageJ software (*45*).

Leaf traits

*Leaf, tissue, and cell dimensions.* Leaf thickness, spongy mesophyll thickness, and spongy mesophyll cell arm diameter, were measured from microCT images of leaf cross-sections. Spongy mesophyll cell arm length, stomata length, and guard cell pair width were measured from paradermal microCT slices. Mean and standard error were calculated from replicates of 15 to 20 measurements from randomly selected slices (leaf thickness, spongy mesophyll thickness) or cells for all species. All measurements were obtained using the FIJI distribution of ImageJ software (*45*).

*Stomatal density and cell packing density.* Stomatal density and cell packing density measurements were measured from microCT images of leaf paradermal slices. Stomatal counts (~10-60 per sample) and spongy mesophyll cell counts (~40-140 per sample) were measured from areas between veins, for direct comparison of these traits in relation to local spongy mesophyll structural organization without the additional factor of displacement of mesophyll area by major and minor veins, which varied between samples and species. All measurements were obtained using the FIJI distribution of ImageJ software (*45*).

*Vein density.* Previously unpublished or otherwise unavailable vein density, $D_v$ (mm mm$^{-2}$), data were contributed by an author (GTR) of this article. Published data sources are listed in References and Notes as reference numbers (*27*, *35*, *46–53*) . All leaf trait data used in our analyses are included in Data Set S1 ("Leaf trait dataset").

*Maximum photosynthetic rate.* To obtain the rate of maximum photosynthesis ($A_{max}$), we combined both empirical and modeled values from the literature as well as our own measurements on plants that were used for microCT imaging. For reported values in the literature, we either used data explicitly stated as maximum photosynthetic rate (sometimes reported as light saturated photosynthesis) directly from tables, or we estimated the maximum photosynthetic rate from light response curves within the light saturated region of the curve. In cases where no literature values were available, we measured maximum photosynthetic rates on leaves using a Li-6400 portable gas exchange system (Licor Inc, Lincoln, NE, USA). Leaves were placed in the chamber and allowed to acclimate to the following conditions until they reached a steady state: PPFD = {0, 50 100, 200, 400, 600, 800, 1000} µmol m$^{-2}$ s$^{-1}$, $CO_2$ = 400 ppm, RH = 37%, VPD = 1.6 kPa. These measurements were performed on 3-4 leaves per species. Published data sources are listed in References and Notes as reference numbers (*54–74*). All $A_{max}$ data used in our analyses are included in Data Set S1 ("Leaf trait dataset").

*Minimum vein spacing distance.* Localized spatial constraints imposed by vein spacing were predicted to influence mesophyll topological pattern. To test this, minimum vein spacing distance was defined as the shortest characteristic distance between veins and measured as the mean length between the highest order veins and neighboring parallel veins or vein segments (Fig. S3). Minimum vein spacing distances were measured from microCT images of leaf paradermal sections replicated 3-10 times per sample using the FIJI distribution of ImageJ software (*45*).

*Spongy mesophyll porosity and surface area.* Leaf image stacks were cropped to spongy mesophyll domains excluding veins, palisade mesophyll, and epidermal layers, and the airspace was segmented by visually and subjectively defining a range of pixel intensity threshold values between a minimum and a maximum grayscale value to optimize airspace classification while minimizing false classification (i.e. non-airspace pixels). The ImageJ plugin BoneJ2 (*75*) was then used to quantify the spongy mesophyll IAS volume, $V_{IAS}$ (µm$^3$), the total spongy mesophyll volume (minus veins), $V_{mes}$ (µm$^3$), and the spongy mesophyll surface area exposed to the IAS, $SA_{mes}$ (µm$^2$) from the cropped, segmented (binary) stacks containing only spongy mesophyll tissue. Mesophyll porosity (m$^3$ m$^{-3}$), was then calculated as the IAS volume as a fraction of the total spongy mesophyll volume. Spongy mesophyll surface area per unit volume, $SA_{mes}/V_{mes}$ (µm$^2$ µm$^{-3}$), was then calculated. Mesophyll surface area per projected leaf area,

$S_m$ ($m^2$ $m^{-2}$), was then calculated as the ratio $SA_{mes}/A_{leaf}$ ($m^2$ $m^{-2}$), where $SA_{leaf}$ ($m^2$) is the surface area of the leaf sample stack, defined as the image width multiplied by the stack depth (i.e. the area of the paradermal view).

*Nearest neighbors.* Nearest neighbors is a classification algorithm used to find the number of edges, i.e. the polygon class, of each spongy mesophyll IAS pore in the paradermal plane, by first determining the center of each pore, then computing a Voronoi diagram, and subsequently counting the number of sides to the Voronoi cell associated with each pore. To pre-process the images, paradermal slices were first made within the spongy mesophyll domain of the microCT leaf image stacks and cropped to exclude veins. The following steps were not carried out for species with spongy mesophyll cells that were disconnected in the paradermal plane, i.e. that did not form a lattice. The airspace was segmented from leaf tissue by visually and subjectively defining a range of pixel intensity threshold values between a minimum and a maximum grayscale value to optimize airspace classification while minimizing false classification (i.e.non-airspace pixels). Edge pixels were then smoothed using the built-in Gaussian Blur filter (0-10 pixel radius of decay). The "watershed irregular features" function for the ImageJ plugin Biovoxxel (*76*) was then used to repair gaps introduced in the tissue structure by the thresholding step, and by cross-referencing the binary image with the grayscale image stack to confirm the fidelity and accuracy of this image processing step. The images were despeckled using the built-in Median filter. The "nearest neighbors" function for the ImageJ plugin Biovoxxel (*76*) was then used to classify pore polygon class using the UEP Voronoi analysis method with pixel size parameters 0-infinity ($mm^2$). Parameters were also set to exclude edge particles from the computation and from the image output. However, even using the edge exclusion parameter, an edge effect was observed where pores at the edges of an image had fewer neighbors compared to pores in the image center. Therefore, the influence of edge effects on the nearest neighbor distribution were analyzed (Fig S4). We found, for images of hexagonal lattices cropped to rectangles with similar side-lengths, that classification accuracy (percent of polygons with 6 neighbors, i.e. "% 6N") decreased sharply below ~150 pores. Using the hexagonal lattice as a proxy for spongy mesophyll, a minimum count of ~150 pores were therefore included in each image for the nearest neighbor analysis, to minimize classification accuracy decreases due to edge effects. Maximum pore counts were limited by veins, oil bodies, and other structures cropped out of the images during image pre-processing that would otherwise locally modify spongy mesophyll tissue organization.

*IAS pore diameter and count.* The following measurements were not carried out for species with spongy mesophyll cells that were disconnected in the paradermal plane, i.e. that did not form a lattice. IAS pore diameter and count were measured from images of paradermally-sliced spongy mesophyll that were either the same images, or images pre-processed following the same method, as in the nearest neighbor analysis. Pore area and count were taken using the ImageJ built-in Particle Analyzer tool. Pore diameter ($D_{pore}$) was then calculated in R (*77*) from pore area ($A_{pore}$) assuming a circular geometry:

$$D_{pore} = \sqrt{\frac{4A_{pore}}{\pi}} \quad (1)$$

Mean and standard deviation were then calculated for each species (Data Set S1).

*Euler characteristics θ, $Z_e$, and $\bar{n}$.* The topological constraints of Euler's Law (*17*) determine the number of vertices (*V*), edges (*E*), and faces (IAS voids, *F*) in a large 2D aggregate according to:

$$F - E + V = 1 \tag{2}$$

so that in a lattice with six edges (E) surrounding each face (F) the number of vertices (V) per face will also be six. This also results in an edge connectivity ($Z_e$) of three, resulting in the familiar hexagonal honeycomb that tessellates 2D space with the least material investment (*22*). The following measurements were not carried out for species with spongy mesophyll cells that were disconnected in the paradermal plane, i.e. that did not form a lattice. To validate the applicability of Euler's Law for 2D slices of spongy mesophyll, first the characteristic internal angle *θ* was measured using the "Inter-trabecular angles" function of the BoneJ2 (*75*) plugin for ImageJ (*45*). This function first thins the image foreground (mesophyll), then creates a graph of the largest skeleton consisting of connection points, or nodes, and connecting edges. The "Use clusters" pruning method was selected to simplify densely clustered nodes. The valence parameter, or edge connectivity $Z_e$, ranged from 3-10, meaning that angles would only be measured at nodes where between 3 and 10 edges met. Inter-edge angle and edge connectivity mean and standard deviation were calculated for each species, and mean and standard deviation were then calculated for the dataset (n = 29). The number of edges per face $\bar{n}$ was calculated from the edge coordination relationship:

$$\bar{n} = \frac{2Z_e}{Z_e - 2} \tag{3}$$

which is a generalized application of Euler's Law (Eqn 1) for a non-regular honeycomb, or to characterize n for a 2D aggregate with varying polygon classes. A Welch two sample t-test was done to compare $\bar{n}$ with the number of nearest neighbors, each of which serves as a proxy for polygon class within the honeycomb lattice. A two-sided analysis ($t(55) = 11.2$, $p < 0.001$) showed there was not a significant difference between $\bar{n}$ (mean = 5.89, s.d. = 0.07) and nearest neighbors (mean = 5.69, s.d. = 0.06).

*Tesselation entropy, Lewis' rule, and Aboav-Weaire law.* Topological methods allow to describe how ordered or regular a structure is. With this in mind, we described dispersion in IAS pore polygon class within the spongy mesophyll using tessellation entropy (Fig. 2H) and by comparison to theoretical predictions from the Aboav-Weaire Law and Lewis' Rule. Tessellation entropy (*S*) describes the degree of order in a system by its polygon class distribution (*23*), where (*n*) is the number of nearest neighbors for each polygon, and *P*(*n*) is the probability of finding *n* nearest neighbors in the system. This was calculated using the nearest neighbors counts for each species according to:

$$S = -\sum_n P(n)\ln[P(n)] \tag{4}$$

with $S = 0$ for a perfectly regular structure. Polygons with fewer than four or greater than nine sides were excluded from the analysis as they typically arise from errors in the segmentation process due to out of plane cell arms that are not included in the nearest neighbors image, thereby creating an artificially large (or small) IAS pore. Mean tessellation entropy and standard deviation was then calculated for all species to which the analysis was applied (n = 29).

The Aboav-Weaire Law (*24*) maintains that any face, i.e. IAS pore, with a lower than average number of edges (n < 6) will introduce a corresponding face with a higher number of edges, so that any five-sided face will have a seven-sided partner within the tessellation, frequently as a neighbor. This is given as:

$$\bar{m} = 5 + \frac{6}{n} \quad (5)$$

where $n$ is the number of edges corresponding to a particular face and $\bar{m}$ is the average number of edges of its $n$ neighbors.

Similarly, Lewis's Law (*25*) relates topological patterns of size dispersion in honeycombs where the area of a given face varies linearly with its number of edges as:

$$\frac{A(n)}{A(\bar{n})} = \frac{n - n_o}{\bar{n} - n_o} \quad (6)$$

where $A(n)$ is the area of a cell with n sides, $A(\bar{n})$ is that of a cell with the average number of sides, and $n_o$ is a constant (Lewis finds $n_o = 2$). On average, this gives larger faces more neighbors.

To calculate the Lewis ratio ($A(n)/A(\bar{n})$) and Aboav-Weaire edge average for each polygon class ($\bar{m}$), the paradermal binary watershed images prepared in the previous step were opened in Fiji and converted to a Voronoi polygon image using the Voronoi module. This created a Voronoi polygon map with IAS polygons in black pixels and the IAS domain boundaries (i.e. the mesophyll cells) in white pixels. These binary Voronoi images were then opened in the Icy image processing software package (*78*) with the EpiTools plugin (*79*). The CellGraph module was used first to create a map of the different IAS polygons while excluding edge domains. These data were then exported using the CellExport module in Icy. This set of scripts and resulting data allowed us to extract the data needed to determine how closely the irregular honeycomb topology of the spongy mesophyll agreed with the predictions of Lewis' Law or the Aboav-Weaire rule. The Aboav-Weiare rule states that cells with more sides than average (i.e. >6) have neighbors with fewer sides than average (i.e. <6), and that introducing a seven sided cell generally requires the introduction of a 5 sided cell often as its neighbor (*17*). Using the EpiTools plugin for Icy, we were able to identify all the IAS polygons within captured in the images, determine the IAS area inscribed by the cell wall boundaries, determine the number of edges of the IAS domains, the number of neighboring IAS polygons, and the area and number of sides of those neighbors. These data were used to then calculate the Lewis ratio and Aboav-Weaire values for each species. Polygons with fewer than four or greater than nine sides were excluded from the analysis as they typically arise from errors in the segmentation process due to out of plane cell arms that are not included in the final Voronoi image, thereby creating an artificially large (or small) IAS pore. Data for each species were then plotted with the model prediction based on Lewis' findings for epithelial cells or the Aboav-Weaire relationship.

Random Forest classification

To determine what variables best predict spongy mesophyll phenotype, Random Forest classification was implemented using the randomForest package (*80*) in the R software environment (*77*). An ensemble of five thousand decision trees with four variables per node were examined, well exceeding the point at which Out-of-Bag (OOB), honeycomb, and non-honeycomb error rate stabilized (~2200 trees). OOB error estimate was 5%, i.e. 95% of samples were correctly classified. The confusion matrix (Table S3) further shows a class error of 0% for the honeycomb phenotype, i.e. all 29/29 species with the honeycomb phenotype were correctly classified. There was a class error of 0.18% for non-honeycomb mesophyll, i.e. 2/11 non-honeycomb species were incorrectly classified as having the honeycomb phenotype. Partial dependence plots show the relative likelihood of classification as either non-honeycomb or honeycomb phenotype according to each variable and are given in Fig. S5. The variable importance plot (Fig. 3A) indicates how important each variable is in classifying the data, with the most important variables at the top of the plot. Importance of variables is measured using mean decrease in accuracy, or the amount by which the model accuracy decreases due to the exclusion of the variable in question.

<u>Power law and linear regression modeling</u>

Allometric relationships were calculated from log-transformed data and linear relationships were calculated from normally distributed, non-transformed data. Model calculations were done using the function "lm" in the R stats package (*77*). For visualization purposes, data were plotted in arithmetic space. In power law figure insets, data were plotted in log-log space for comparison.

To determine the evolutionary coordination between traits, we constructed a phylogeny from the list of taxa using Phylomatic (v. 3) and its stored family-level supertree (v. R20120829) using the R package brranching (*81*). Following published methods (*32*), we compiled node ages of named crown group from fossil-calibrated estimates of crown group ages (*82–84*). Of the 32 internal nodes in our phylogeny, 31 of them had published ages, which were assigned to nodes and branch lengths between dated nodes smoothed using the function 'bladj' in the software Phylocom (v. 4.2) (*85*).

We tested whether there was correlated evolution between traits (Table S2) using phylogenetic least squares regression with a Brownian motion correlation structure, using the R packages nlme (*86*) and ape (*87*). For analyses of correlated trait evolution, traits were log-transformed to improve normality prior to regression analyses.

<u>Surface area and volume calculations of idealized isodiametric and triply-lobed cells</u>

Surface areas of isodiametric (spherical) and triply-lobed cells with the same volume were approximated using idealized 3D geometrical representations. Isodiametric cells were modeled as spheres (Fig. S6A), and triply-lobed cells were modeled as three cylinders connected to a triangular prism with an equilateral base (Fig. S6B). Volume of a triply-lobed cell ($V_{arm\ cell}$) was first calculated according to:

$$V_{arm\,cell} = \frac{\sqrt{3}}{4}A_D^3 + 3[\pi\left(\frac{A_D}{2}\right)^2(A_L - \frac{1}{2}\sqrt{\left(\frac{A_D}{2}\right)^2 + (A_D)^2})] \quad (7)$$

with cell diameter $A_D$ and cell arm length $A_L$. Surface area of the arm cell ($SA_{arm\ cell}$) included the upper and lower faces of the triangular prism and the side and (one) base of each cylinder:

$$SA_{arm\ cell} = \frac{\sqrt{3}}{2}A_D^2 + 3[(\pi A_D)(A_L - \frac{1}{2}\sqrt{\left(\frac{A_D}{2}\right)^2 + (A_D)^2}) + \pi\left(\frac{A_D}{2}\right)^2] \tag{8}$$

Volume was considered as conserved between the arm cell and isodiametric cell:

$$V_{isodiametric\ cell} = V_{arm\ cell} \tag{9}$$

and surface area of the isodiametric cell ($SA_{isodiametric\ cell}$) was then calculated as:

$$SA_{isodiametric\ cell} = 4\pi r^2 \tag{10}$$

with the isodiametric cell radius (r) given by:

$$r = \sqrt[3]{\frac{3V_{isodiametric\ cell}}{4\pi}} \tag{11}$$

Surface area to volume ratios were then calculated for each cell type using the measured $A_D$ and $A_L$ values. Power law regressions were calculated from log-transformed data and analyzed using the function "lm" in the R stats package (*77*). For visualization purposes, data were plotted in arithmetic space. In figure insets, data were plotted in log-log space for comparison.

**Supplementary Figures**

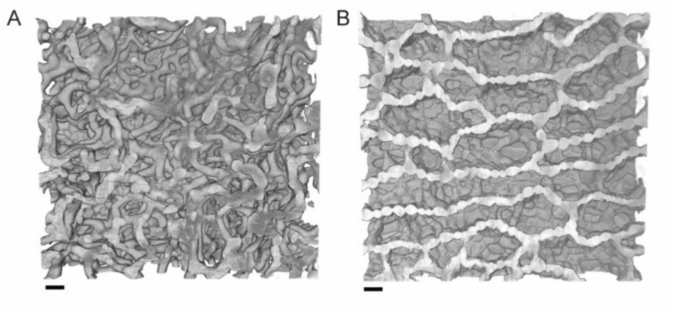


**Fig. S1. Spongy mesophyll structural outliers found in species with reticulate venation. (A)** Paradermal microCT images of the spongy mesophyll of spinach (*Spinacia oleracea*), with elongated cell arms forming large, convoluted and continuous airspaces. **(B)** Paradermal microCT image of the spongy mesophyll of water lily (*Nuphar polysepala*), where multiple layers of the spongy mesophyll formed a lattice-like structure as found in species within the honeycomb class, but arranged such that the IAS domains were elongated to form a highly porous tissue layer, presumably to increase buoyancy. Bars = 50 μm.

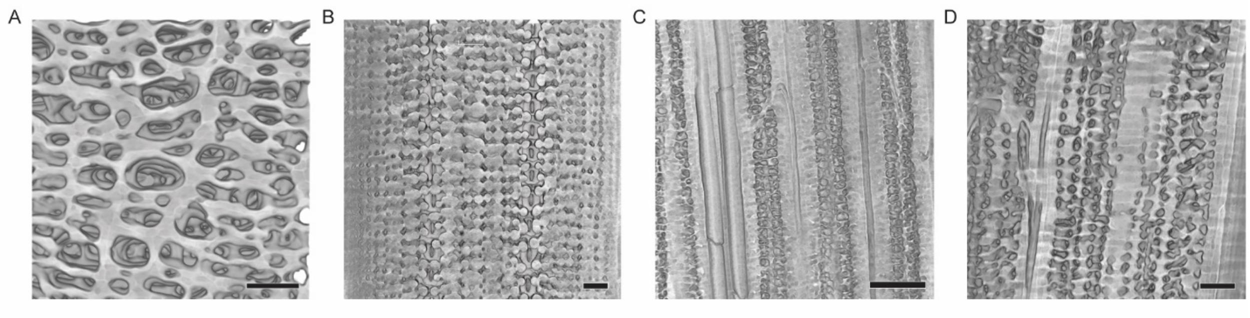


**Fig. S2. Structural variation in spongy mesophyll in leaves with parallel venation. (A)** Paradermal microCT image of Cycas revoluta with spongy mesophyll cells arranged in an orthogonal lattice. (B) Paradermal microCT image of *Pinus monitcola* where the spongy mesophyll adjacent to the stomatal guard cells are arranged in an orthogonal lattice. (C) Paradermal microCT image of the spongy mesophyll in rice (Oryza sativa) with parallel venation. (D) Paradermal microCT image of the spongy mesophyll between the parallel veins of *Calamagrostis arundinacea*. Bars = 50 µm.

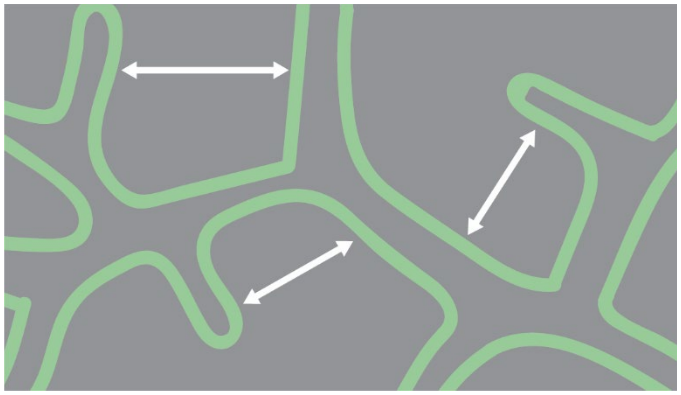

**Fig. S3. Diagram showing minimum vein spacing measurements.** Measurements are made between the highest order vein and neighboring parallel vein.

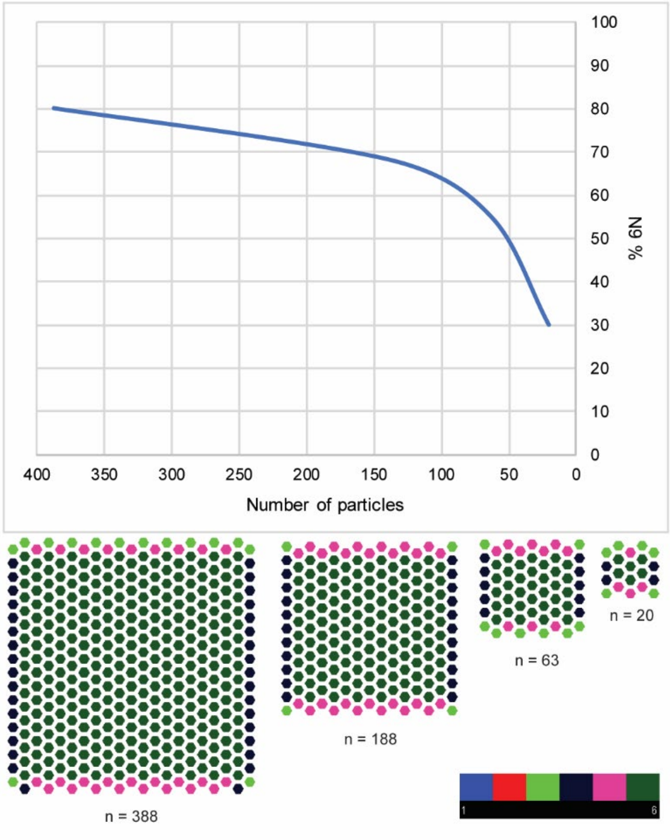


**Fig. S4. Nearest neighbors edge effect analysis.** Influence of edge effects on model accuracy for a range of aggregate sizes.

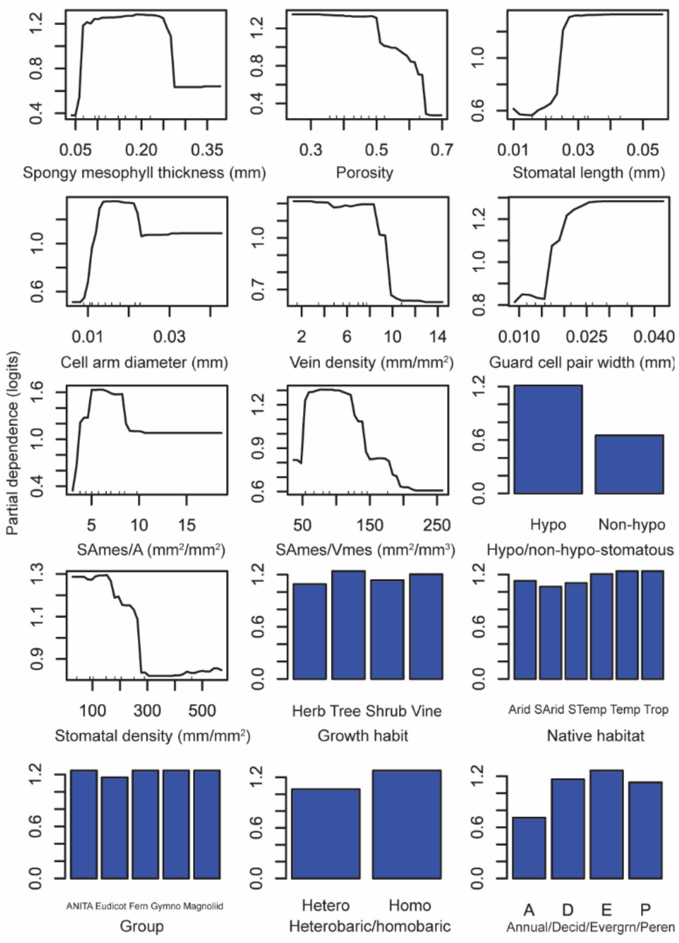


**Fig. S5. Partial dependence plots for random forest predictors.** Partial dependence (logits) for model predictors (mean accuracy decrease > 0; predictors with three highest importance rankings shown in Fig. 3B-D).

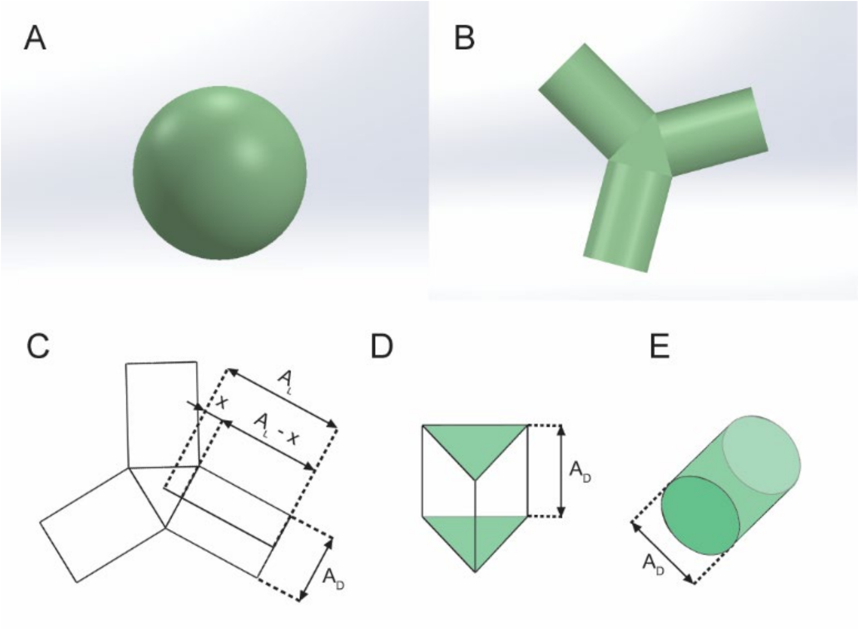


**Fig. S6. Schematic diagram of idealized cell geometries used in surface area and volume calculations for individual cells. (A)** Sphere. **(B)** Triply-armed cell. **(C)** Top view of triply-armed cell. **(D)** Prismatic center of triply-armed cell. **(E)** Cylindrical arm of triply-armed cell.

### Supplementary Tables

| | | 1 | 2 | 3 | 4 | 5 | 6 | 7 | 8 | 9 | 10 | 11 | 12 | 13 |
|---|---|---|---|---|---|---|---|---|---|---|---|---|---|---|
| 1 | Arm diameter | | | | | | | | | | | | | |
| 2 | Arm length | **0.95**[a] | | | | | | | | | | | | |
| 3 | Leaf thickness | **0.75**[a] | **0.64**[a] | | | | | | | | | | | |
| 4 | Spongy thickness | **0.81**[a] | **0.80**[a] | **0.85**[a] | | | | | | | | | | |
| 5 | Vein density | **-0.63**[a] | **-0.71**[a] | **-0.42**[b] | **-0.57**[a] | | | | | | | | | |
| 6 | Minimum vein spacing | **0.79**[a] | **0.82**[a] | **0.40**[c] | **0.60**[a] | **-0.64**[c] | | | | | | | | |
| 7 | Stomata length | **0.79**[a] | **0.78**[a] | **0.46**[b] | **0.60**[a] | **-0.63**[a] | **0.76**[a] | | | | | | | |
| 8 | Stomatal density | **-0.69**[a] | **-0.72**[a] | **-0.37**[c] | **-0.54**[a] | **0.75**[a] | **-0.75**[a] | **-0.76**[a] | | | | | | |
| 9 | $SA_{mes}/V_{mes}$ | **-0.94**[a] | **-0.94**[a] | **-0.77**[a] | **-0.86**[a] | **0.61**[a] | **-0.72**[a] | **-0.69**[a] | **0.61**[a] | | | | | |
| 10 | $SA_{mes}/A$ | 0.02 | -0.06 | 0.10 | 0.08 | -0.07 | -0.08 | 0.05 | 0.00 | 0.05 | | | | |
| 11 | Porosity | -0.17 | -0.11 | -0.01 | -0.06 | 0.19 | **-0.29**[c] | -0.10 | 0.13 | -0.04 | -0.15 | | | |
| 12 | Cell packing density | **-0.91**[a] | **-0.97**[a] | **-0.58**[a] | **-0.75**[a] | **0.72**[a] | **-0.86**[a] | **-0.79**[a] | **0.72**[a] | **0.89**[a] | 0.15 | 0.15 | | |
| 13 | IAS pore diameter (n = 29) | **0.75**[a] | **0.83**[a] | **0.59**[a] | **0.65**[a] | **-0.46**[c] | **0.32**[c] | **0.66**[a] | **-0.37**[c] | **-0.85**[a] | 0.09 | **0.37**[c] | **-0.80**[a] | |
| 14 | $A_{max}$ ( n = 29) | **-0.63**[a] | **-0.67**[a] | **-0.59**[a] | **-0.71**[a] | **0.51**[b] | **-0.52**[b] | **-0.41**[c] | **0.45**[c] | **0.72**[a] | 0.12 | 0.16 | **0.60**[a] | **-0.48**[c†] |

[a]$p < 0.001$

[b]$0.001 \leq p < 0.01$

[c]$0.01 \leq p < 0.05$

[†]n = 23

**Statistically significant,** statistically non-significant

**Table S1. Spearman correlation matrix for continuous leaf trait variables (n = 40).**

| | | 1 | 2 | 3 | 4 | 5 | 6 | 7 | 8 | 9 | 10 | 11 | 12 | 13 |
|---|---|---|---|---|---|---|---|---|---|---|---|---|---|---|
| 1 | Arm diameter | | | | | | | | | | | | | |
| 2 | Arm length | **37, 9.6, <0.0001** | | | | | | | | | | | | |
| 3 | Leaf thickness | **37, 4.5, 0.0001** | **37, 3.0, 0.0045** | | | | | | | | | | | |
| 4 | Spongy thickness | **37, 5.8, <0.0001** | **37, 5.5, <0.0001** | **37, 11.2, <0.0001** | | | | | | | | | | |
| 5 | Vein density | 37, -2.0, 0.055 | **37, -3.4, 0.0017** | 37, -1.3, 0.2111 | **37, -2.4, 0.0209** | | | | | | | | | |
| 6 | Minimum vein spacing | **37, 4.2, 0.0002** | **37, 6.7, <0.0001** | 37, -0.1, 0.9380 | 37, 1.5, 0.1524 | **37, -3.6, 0.0009** | | | | | | | | |
| 7 | Stomata length | **37, 4.7, <0.0001** | **37, 4.4, 0.0001** | 37, 1.7, 0.0999 | **37, 2.9, 0.0062** | **37, -3.2, 0.0029** | **37, 3.9, 0.0004** | | | | | | | |
| 8 | Stomatal density | **37, -2.4, 0.02** | **37, -3.0, 0.0051** | 37, 0.3, 0.7492 | 37, -1.0, 0.3193 | **37, 2.7, 0.0115** | **37, -4.5, 0.0001** | **37, -5.8, <0.0001** | | | | | | |
| 9 | $SA_{mes}/V_{mes}$ | **37, -13.1, <0.0001** | **37, -11.1, <0.0001** | **37, -5.8, <0.0001** | **37, -8.23, <0.0001** | **37, 2.6, 0.0145** | **37, -3.3, 0.0020** | **37, -4.1, 0.0002** | 37, 1.9, 0.0722 | | | | | |
| 10 | $SA_{mes}/A$ | 37, 1.01, 0.3158 | 37, -0.6, 0.5777 | 37, 1.9, 0.0656 | 37, 1.6, 0.1121 | 37, 0.1, 0.9451 | 37, -1.1, 0.2793 | 37, 0.7, 0.4907 | 37, 0.8, 0.4394 | 37, -0.3, 0.7327 | | | | |
| 11 | Porosity | 37, -0.33, 0.7427 | 37, 0.37, 0.7117 | 37, -0.3, 0.7875 | 37, -0.1, 0.9432 | 37, -0.2, 0.7750 | 37, -0.1, 0.9368 | 37, 1.0, 0.3297 | 37, -1.1, 0.2681 | 37, -0.7, 0.4949 | 37, -1.2, 0.2237 | | | |
| 12 | Cell packing density | **37, -7.1, <0.0001** | **37, -14.5, <0.0001** | **37, -2.3, 0.0278** | **37, -4.0, 0.0003** | **37, 4.0, 0.0003** | **37, -7.4, <0.0001** | **37, -4.5, <0.0001** | **37, 2.6, 0.0128** | **37, 7.5, <0.0001** | 37, 0.6, 0.5491 | 37, -0.9, 0.3988 | | |
| 13 | IAS pore diameter | **26, 3.6, 0.0016** | **26, 6.5, <0.0001** | **26, 2.4, 0.0258** | **26, 3.1, 0.0055** | 26, -1.5, 0.1569 | 26, 1.2, 0.2519 | **26, 4.5, 0.0001** | 26, -1.5, 0.1322 | **26, -5.6, <0.0001** | 26, 0.1, 0.9185 | **26, 3.4, 0.0024** | **26, -4.7, 0.0001** | |
| 14 | $A_{max}$ | 27, -1.9, 0.0687 | **27, -3.0, 0.0057** | 27, -1.0, 0.3459 | **27, -0.2, 0.0476** | 27, 1.6, 0.1323 | 27, -1.5, 0.1555 | 27, -0.9, 0.3774 | 27, 1.2, 0.2442 | **27, 2.6, 0.0138** | 27, 0.9, 0.3755 | 27, -0.1, 0.9421 | 27, 1.9, 0.0662 | 21, -0.8, 0.4604 |

Degrees of freedom, t-values, and p-values reported in each cell.

**Statistically significant, s**tatistically non-significant

**Table S2. Degrees of freedom, t-values, and p-values for pairwise generalized linear models taking into account phylogenetic non-independence between data points.**

| | Honeycomb | Non-Honeycomb | Class Error |
|---|---|---|---|
| Honeycomb | 29 | 0 | 0 |
| Non-Honeycomb | 2 | 9 | 0.18 |

**Table S3. Confusion matrix for random forest classification of spongy mesophyll phenotype.**

**Movie S1.**

MicroCT volume rendering of a coffee leaf (*Coffea arabica*) with a paradermal bisection, opening to reveal the honeycomb structure of the spongy mesophyll. Tissue dimensions are approximately 50 µm across the horizontal edge.

**Movie S2.**

MicroCT volume rendering of a star anise leaf (*Illicium floridanum*) with a paradermal bisection, opening to reveal the honeycomb structure of the spongy mesophyll. Tissue dimensions are approximately 50 µm across the horizontal edge.
